# Analysis of polygenic selection in purebred and crossbred pig genomes using Generation Proxy Selection Mapping

**DOI:** 10.1101/2022.09.19.508567

**Authors:** Caleb J. Grohmann, Caleb M. Shull, Tamar E. Crum, Clint Schwab, Timothy J. Safranski, Jared E. Decker

## Abstract

**Background:** Artificial selection on quantitative traits using breeding values and selection indices in commercial livestock breeding populations causes changes in allele frequency over time, termed polygenic selection, at causal loci and the surrounding genomic regions. Researchers and managers of pig breeding programs are motivated to understand the genetic basis of phenotypic diversity across genetic lines, breeds, and populations using selection mapping analyses. Here, we applied Generation Proxy Selection Mapping (**GPSM**), a genome-wide association analysis of SNP genotype (38,294 to 46,458 SNPs) of birth date, in four pig populations (15,457, 15,772, 16,595 and 8,447 pigs per population) to identify loci responding to artificial selection over a span of five to ten years. Gene-drop simulation analyses were conducted to validate GPSM results. Selection signatures within and across each population of pigs were compared in the context of commercial pork production.

**Results:** Forty-nine to 854 loci were identified by GPSM as under selection (*Q*-values less than 0.10) across 15 subsets of pigs based on population combinations. The number of significant associations increased as populations of pigs were pooled. In addition, several significant associations were identified in more than one population. These results indicate concurrent selection objectives, similar genetic architectures, and shared causal variants responding to selection across populations. Negligible error rates (less than or equal to 0.02%) of false-positive associations were identified when testing GPSM on gene-drop simulated genotypes, suggesting that GPSM distinguishes selection from random genetic drift in actual pig populations.

**Conclusions:** This work confirms the efficacy and accuracy of the GPSM method in detecting selected loci in commercial pig populations. Our results suggest shared selection objectives and genetic architectures across swine populations. Identified polygenic selection highlights loci important to swine production.

## Background

Artificial selection in pigs, over the past 300 years, has led to the formation of pig breeds with well-defined breed characteristics and considerable across breed variation in phenotypes related to economically relevant traits [1]. Pig breeders placing selection pressure on certain qualitative phenotypes such as coat color and ear morphology and quantitative phenotypes such as feed efficiency, average daily gain, and backfat depth has left signatures of selection, known as selective sweeps, across the genomes of pig populations [2]. In general, intense signatures of selection, which are large, rapid changes in allele frequency which drag neighboring variation, are associated with phenotypes that underly the divergence of pig breeds, and these “hard selective sweeps” have been identified in pig genomes by several studies [1–3]. However, pig breeders, since the adoption of mixed model methods in animal breeding in the 1980s, have been more concerned with selection for increased rates of genetic gain in quantitative traits [4]. The selection index has served as the most efficient method in assessing and improving the aggregate genetic merit of pigs by combining data from multiple traits in a breeding objective [5,6]. Economically relevant traits that are utilized in selection indices are generally quantitative in nature; thus, these traits are controlled by large numbers of causal variants, termed quantitative trait loci (**QTL**) first by Geldermann [7,8] in 1975. Artificial selection using selection indices in pig breeding programs has been proven to cause significant changes to the mean phenotype of any one trait considered within the breeding objective [9–11]. However, artificial selection pressure, especially over relatively short time scales, causes only subtle changes to allele frequencies at QTLs across the genome [12,13]. While allele frequencies change at QTLs for traits included in the selection index, neighboring loci to these QTLs can also “hitchhike” along with the primary locus under selection due to linkage disequilibrium [14]. In addition, allele frequencies at loci that affect traits that are not explicitly included in the selection index have been shown to undergo frequency changes as a result of artificial selection pressure applied in livestock breeding programs [13,14].

There is much interest within the area of livestock genomics in deciphering the genetic basis of phenotypic diversity in species raised in animal agriculture for meat production [13,14]. Understanding selection in livestock populations is of paramount importance when evaluating the genomic basis of phenotypic variation within a genetic line, breed, or entire livestock population over time. The identification of selection detects loci that have been subjected to consistent increases or decreases in allele frequency of greater intensity than random genetic drift [15–17]. Unlike hard or soft sweeps, polygenic selection does not leave distinctive signatures on the genome [13]. With current technologies such as single nucleotide polymorphism (**SNP**) arrays, temporally distributed genotypes, and increased computing resources, statistical analysis of polygenic selection is efficient. Identification of regions of the genome that have been altered due to artificial selection pressure is highly beneficial in ascertaining QTLs under selection [13]. When results of selection mapping analyses are combined with results from phenotype-based genome wide association analyses (**GWAAs**), QTLs associated with phenotypic variation of traits within breeding objectives can be confirmed [18]. Moreover, there are opportunities within selection mapping analyses to evaluate results within or across genetic lines or breeds, which can highlight differences in selection objectives across livestock breeding programs. Selection mapping analyses are not limited to increasing knowledge with respect to selection and evolution of species. Further, using results from selection analyses, SNP assays used for genomic prediction of breeding values in livestock populations can be refined in order to reduce extraneous statistical noise and increase prediction accuracy. This feature selection of SNPs can be accomplished by excluding SNPs that have not undergone significant changes due to selection or have not contributed to phenotypic variation in traits in the breeding objective.

Generation Proxy Selection Mapping (**GPSM**) has been used as an analytical method for detection of polygenic selection loci in populations [13,14,19]. In this approach, animal birth date (or other generation proxy) is fit as the dependent variable, and SNPs that are strongly associated with or predictive of birth date are identified. If a SNP is under directional selection pressure, changes in allele frequency will generally be consistent over time, and an animal’s genotype will be strongly predictive of birth date [13,14]. In addition, a major advantage in the applicability of GPSM methodology to livestock species over other methods, such as site frequency spectrum and linkage disequilibrium-based methods, is the ability to adjust for demography and confounding due to non-random ascertainment of genotype samples, population structure, inbreeding, or kinship with the use of a genomic relationship matrix (**GRM**) [13,14]. Generation proxy selection mapping has been proven effective and accurate in identifying loci with changes in allele frequency due to polygenic selection (as opposed to loci-specific allele frequency changes due to random genetic drift) in beef cattle populations that have been exposed to artificial selection for approximately 50 years [13]. However, there are stark differences between cattle breeding programs and swine breeding programs. For example, generation intervals in pigs are much shorter than in cattle (2 to 2.5 versus 4 to 5 years, respectively) [20]. Thus, in traits with similar heritability and assuming similar selection intensity, comparable amounts of genetic gain are expected in approximately half the time for pig populations versus the cattle populations. Moreover, due to increasing adoption of specialized sire and dam lines, the classical “breeding pyramid”, and vertical integration in the swine industry, breeding objectives within a population of pigs tend to be more focused than breeding objectives within cattle breeds, where each breeder and farm have their own breeding objectives that may be poorly defined. The described differences between cattle and swine breeding programs contribute to variation in the effect of artificial selection on allele frequencies over time. The objectives of the current study were to 1) use GPSM to identify loci under artificial selection in three purebred populations and one crossbred population of pigs and 2) compare and contrast the effect of artificial selection patterns among genotypes of each population in the context of commercial pig production.

## Methods

### Population background

Four populations of pigs were used in the present study (data owned by The Maschhoff’s, LLC, Carlyle, Illinois, USA). Within each population, a selection index was utilized to identify boars and gilts with superior genetic merit to return to the breeding population at the nucleus level. Breeding population-specific selection indices for all populations included expected progeny differences (**EPDs**) for growth and carcass traits such as increased feed efficiency and average daily gain, decreased backfat depth, and increased *Longissimus* muscle area. In addition, selection indices for two of the four breeding populations (Landrace and Yorkshire) also emphasized maternal reproductive traits and included EPDs for increased number and weight of piglets born and weaned.

### Pedigree and genotype data

A pedigree consisting of individual, sire, and dam identification, birth date, and genetic line for 1,247,982 pigs was provided by The Maschhoff’s. Within the entire pedigree, there were, 117,836, 242,699, 216,915, and 690,202 pigs from the Duroc, Landrace, Yorkshire, and Crossbred populations, respectively. Summary information regarding the number of sires and dams, founder pigs, and generations within each population is presented in Table 1. Complete sire and dam pedigree information for individuals dating back to founder pigs was present for the purebred populations. Due to the nature of the commercial testing program, sire pedigree information dating back to Duroc founder pigs was known for the Crossbred pigs; however, only the identification numbers of the dams of these Crossbred pigs were provided. On a subset of 16,802, 19,342, 18,368, and 8,532 pigs from the Duroc, Landrace, Yorkshire, and Crossbred populations, respectively, SNP assays were collected using a GGP Porcine 50K (Neogen, Corp., Lansing, Michigan, USA) genotyping array. Genomic coordinates for each SNP were from the Sscrofa 10.2 reference genome [21]. Sample collection and subsequent genotyping was conducted on all viable male selection candidates prior to removal from performance testing trials. In addition, all female animals selected to return to the nucleus breeding herd were genotyped. Summary information regarding the number of sires and dams, founder pigs, birth date ranges, and generations for genotyped pigs within each population is provided in Table 2.

**Table 1.**
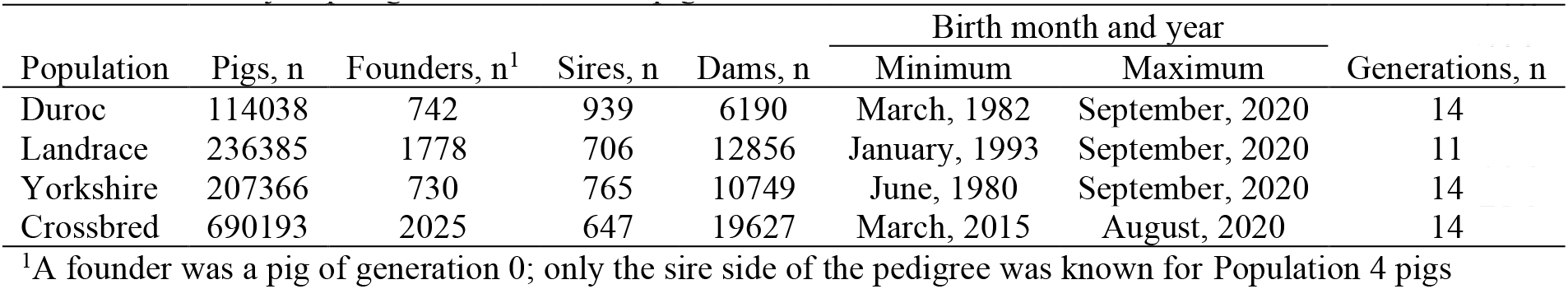
Summary of pedigree records for all pigs.

| Population | Pigs, n | Founders, n<sup>1</sup> | Sires, n | Dams, n | Birth month and year |  | Generations, n |
| --- | --- | --- | --- | --- | --- | --- | --- |
|  |  |  |  |  | Minimum | Maximum |  |
| Duroc | 114038 | 742 | 939 | 6190 | March, 1982 | September, 2020 | 14 |
| Landrace | 236385 | 1778 | 706 | 12856 | January, 1993 | September, 2020 | 11 |
| Yorkshire | 207366 | 730 | 765 | 10749 | June, 1980 | September, 2020 | 14 |
| Crossbred | 690193 | 2025 | 647 | 19627 | March, 2015 | August, 2020 | 14 |

<sup>1</sup>A founder was a pig of generation 0; only the sire side of the pedigree was known for Population 4 pigs

**Table 2.**
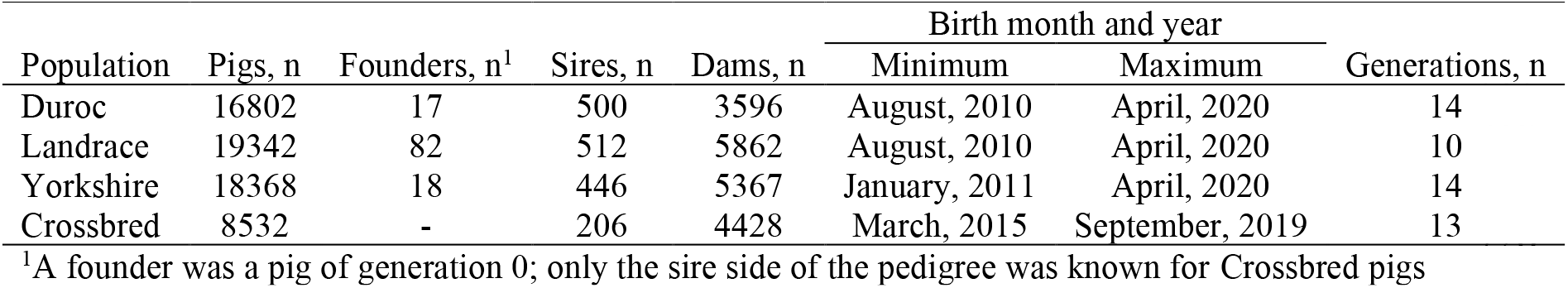
Summary of pedigree records for all genotyped pigs.

| Population | Pigs, n | Founders, n<sup>1</sup> | Sires, n | Dams, n | Birth month and year |  | Generations, n |
| --- | --- | --- | --- | --- | --- | --- | --- |
|  |  |  |  |  | Minimum | Maximum |  |
| Duroc | 16802 | 17 | 500 | 3596 | August, 2010 | April, 2020 | 14 |
| Landrace | 19342 | 82 | 512 | 5862 | August, 2010 | April, 2020 | 10 |
| Yorkshire | 18368 | 18 | 446 | 5367 | January, 2011 | April, 2020 | 14 |
| Crossbred | 8532 | - | 206 | 4428 | March, 2015 | September, 2019 | 13 |

<sup>1</sup>A founder was a pig of generation 0; only the sire side of the pedigree was known for Crossbred pigs

### Preparation of genotype data and overview of analyses

The dependent variable for all analyses was birth date (**AGE**) calculated as the difference, in months, between each pig’s birth month and January 2006. Pigs from the entire dataset of genotyped pigs were separated into 15 subsets based on population or combination of populations. Analyses were conducted using only SNPs located on the autosomal chromosomes for *Sus scrofa*, which were chromosomes 1 through 18. Genotype quality control was performed in PLINK v1.9 [22] for each subset. Any SNP with a genotype call rate less than 0.90 or a minor allele frequency less than 0.01 was filtered from the data. In addition, individual pigs that had a genotype call rate less than 0.90 were filtered from the dataset.

Percent Duroc, Landrace, and Yorkshire ancestry was predicted for each pig using fastSTRUCTURE [23], with the *K* parameter set to 3. Purebred pigs that were predicted to be less than 95% of their assigned genetic line (Duroc, Landrace, or Yorkshire) were removed from all subsequent analyses, as these may represent sample swaps or other pedigree errors. While predicted breed proportions were estimated for the Crossbred pigs, none were removed from the genotyped sample, as deviations from expected breed proportions cannot be distinguished from deviations due to Mendelian sampling. Genomic relationship matrices (**GRMs**) were estimated for each subset using the software GCTA v1.93.2 [24] and the method described by Yang et al. [25], and these GRMs were utilized in all subsequent analyses. To visualize the genomic relatedness between lines, the ‘pca’ function of GCTA [26] was also used to conduct a principal component analysis (**PCA**) on a GRM for all Duroc, Landrace, Yorkshire, and Crossbred pigs. A summary of the number of pigs and SNPs after quality control and all subsequent analyses performed for each subset is presented in Table 3. Descriptive statistics of AGE by genetic line were calculated using the ‘dplyr’ package [27] of the statistical analysis software R [28]. Figures were generated using the ‘ggplot2’ [29] package of R.

**Table 3.**
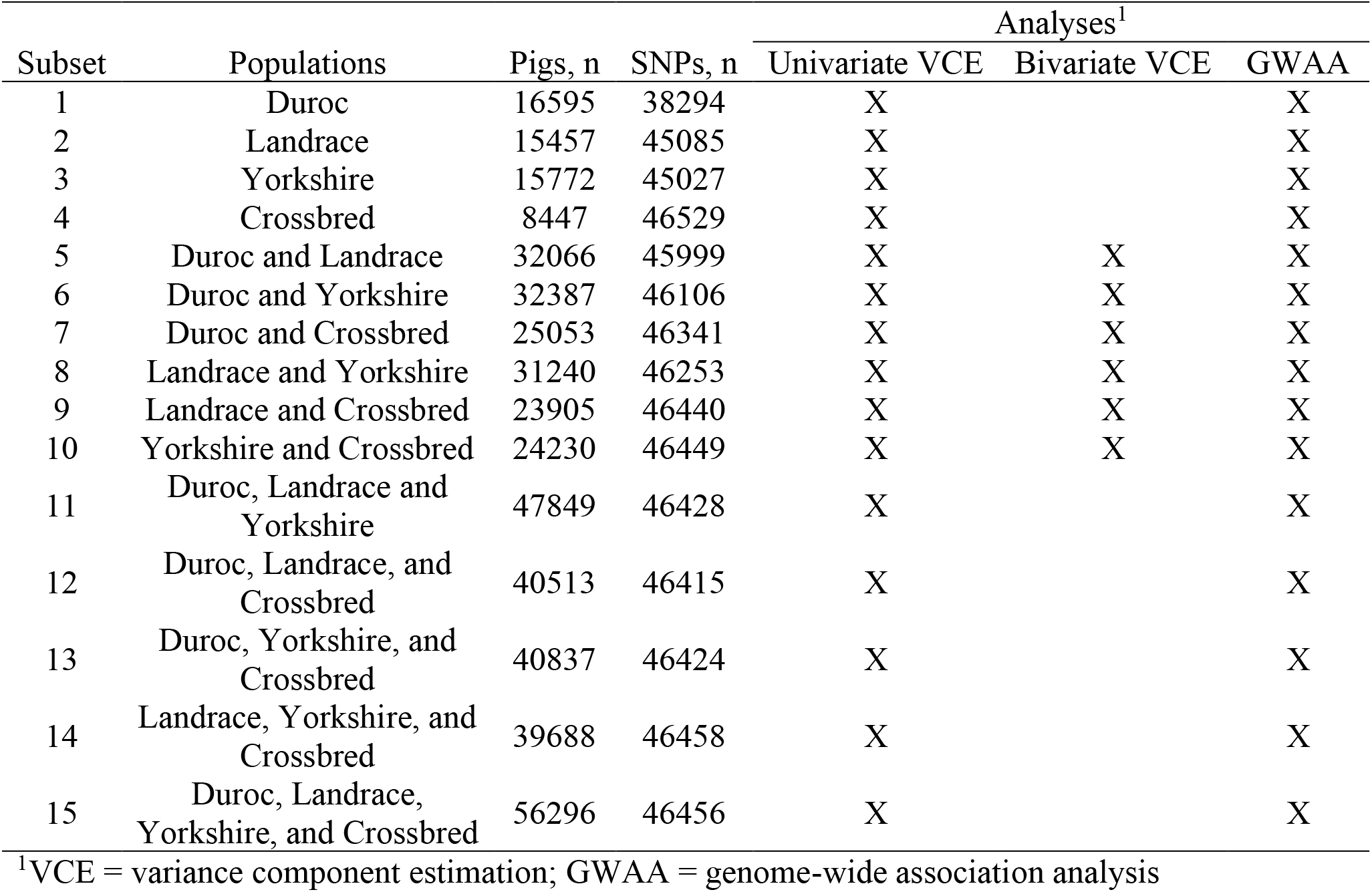
Summary of subsets of genotyped pigs and conducted analyses after genotype quality control.

Depending on data subset, certain combinations of the following three statistical analyses were performed on AGE: 1) univariate variance component estimation, 2) bivariate variance component estimation, and 3) univariate genome-wide association using a mixed linear model to estimate SNP effects.

### Univariate variance component estimation

To estimate the proportion of variance in AGE explained by genome-wide SNPs (**PVE**; also termed SNP heritability) for each subset (Table 3), the following model was fit using GCTA:

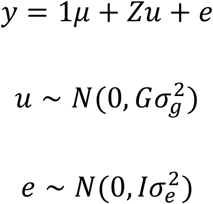

where **y** is a vector of observations for AGE, μ is the overall mean for AGE, **u** is a vector of random polygenic effects, **Z** is an incidence matrix relating AGE in **y** to random polygenic effects in **g**, and **e** is a vector of random residuals, **G** is the genomic relationship matrix, and **I** is an identity matrix. Additive genetic 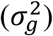 and residual 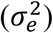 variance components were derived using average information restricted maximum likelihood. The PVE was then estimated as follows:

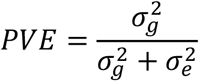

### Bivariate variance component estimation

Genetic correlations (**rG**) between each population (Table 3) for AGE were estimated using bivariate mixed linear models, fitted in GCTA, of the following form:

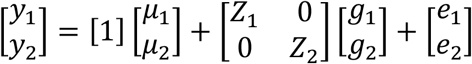

where ***y***_**1**_ and ***y***_**2**_ are vectors of observations for AGE for the two populations, *µ*_1_ and *µ*_2_ are the overall means for AGE for each population, respectively, ***g***_**1**_ and ***g***_**2**_ are vectors of random polygenic effects for each pig in the two populations, ***e***_**1**_ and ***e***_**2**_ are residuals for AGE of the two populations, and ***Z***_**1**_ and ***Z***_**2**_ are incidence matrices for the random polygenic effects in ***g***_**1**_ and ***g***_**2**_, respectively. Additive genetic variance of ***g***_**1**_ and ***g***_**2**_ (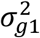 and 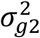, respectively), additive genetic covariance between ***g***_**1**_ and ***g***_**2**_ (*σ*_*g*1,*g*2_), and residual variance of ***e***_**1**_ and ***e***_**2**_ (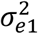 and 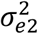, respectively) were estimated using average information restricted maximum likelihood with the variance-covariance matrix (**V**) defined as:

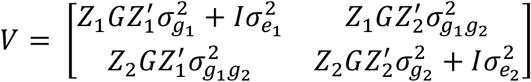

where **G** and **I** were the genomic relationship and identity matrix, respectively. Genetic correlations were then estimated by GCTA using the following formula:

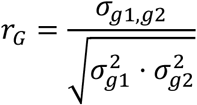

### Generation Proxy Selection Mapping (GPSM)

Generation Proxy Selection Mapping analyses were conducted to detect SNPs with allele frequency changes over time within each subset (Table 3). To accomplish this, univariate mixed linear models were fit in GCTA as part of genome-wide association analyses of AGE, with the models defined as follows:

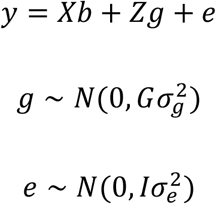

where **y** is the pig’s generation proxy (AGE), and **X** is an incidence matrix that relates SNPs to AGE for each pig and **b** is the estimated allele substitution effect for each SNP. Confounding due to population structure, relatedness, and inbreeding are controlled by the random polygenic term **g**, and **Z** is an incidence matrix for the effect **g**. In addition, **G** is the genomic relationship matrix, and **I** is an identity matrix. Additive genetic 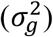 and residual 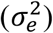 variance components were estimated using average information restricted maximum likelihood. However, these variance components were not of interest as a part of the GPSM analyses as they were estimated previously as a part of the univariate variance component estimation analysis. *P*-values of estimated SNP effects were converted to false discovery rate (**FDR**) corrected *Q*-values using the ‘qvalue’ package [30] of R, and a significance threshold of *Q* < 0.10 was used for all analyses.

### Variance component and GPSM analyses using simulated data

Variance component and GPSM analyses of purebred pigs (subsets 1, 2, 3, 5, 6, 8, and 11; Table 3) were conducted on gene-drop simulated genotype data produced using the package ‘AlphaSimR’ [31] with the pedigrees of the analyzed pigs. The objective of these gene-drop analyses was to ensure GPSM results performed on real data were due to artificial selection as opposed to random genetic drift. For the univariate variance component estimation and GPSM analyses for each of the subsets 1 through 3, 5, 6, 8, and 11 (Table 3), 5,000 founder pig haplotypes were simulated using AlphaSimR’s MaCS [32] wrapper. Each of the simulated haplotypes contained 90,000 segregating sites located evenly across 18 chromosomes. Then, using the pedigreeCross function [31], founder pigs in the pedigree of each subset were assigned genotypes at random from the simulated population of 5,000 pigs. Simulated founder pig genotypes were then dropped through each pedigree to simulate the exact matings that have occurred in The Maschhoff’s breeding program (allele inherited by progeny was randomly assigned). Lastly, pigs with genotypes used in the real analyses were pulled from each subset along with a “SNP chip” of randomly selected loci equivalent to the number of SNPs used in the real analyses (Table 3). Univariate variance component estimation and GPSM analyses were conducted using the same statistical models and software, the simulated genotypes, and the AGE values from the real analyses.

In bivariate variance component analyses on simulated data (subsets 5, 6, and 8; Table 3), founder pig haplotypes were simulated two different ways. First, founder pig haplotypes were simulated as one group that consisted of 15,000 founder pigs (Method 1). The objective of this method was to simulate a scenario where each combination of populations had recently diverged; thus, the founder animals for each population have the same genotypes. For the second method, founder pig haplotypes were simulated separately for each population, the random number generator in R was changed between each simulation, and then the two founder pig haplotypes were combined (Method 2). Using Method 2, the simulated genotypes were vastly different between founder pigs in each population combination, which represented pairs of populations that were completely unrelated. Samples of pigs with simulated genotypes were created in the same manner as described above for the univariate analyses. Bivariate variance component analyses were then conducted using both samples of simulated genotypes from each method and each pairwise comparison of subsets 5, 6, and 8. Results from all analyses using simulated data and analyses using real data were then compared in a one-to-one fashion.

### Investigation of GPSM associations

The number of shared significant GPSM associations were visualized using the R package ‘UpSetR’ [31]. The ‘GALLO’ package of R [43] was used to identify positional candidate genes.

## Results

### Descriptive statistics and principal components analysis

Descriptive statistics of AGE for each subset are presented in Table 4. In addition, raw distributions of AGE are shown in Figure 1 for all pigs (subset 15; Table 3) and Figure 2 for subsets 1 through 4 (Table 3). The histograms of AGE depict the frequency of genotype sampling across all populations and within each population for the duration of The Maschhoff’s breeding program (Figures 1 and 2, respectively). In general, descriptive statistics for AGE were similar across each subset (Table 4). However, the range and standard deviation of AGE for the Crossbred pigs was less than that of the other subsets, as genotyping for these pigs did not begin until March of 2015 (Table 2). Thus, the number of genotyped Crossbred pigs was approximately half of the number of Duroc, Landrace, and Yorkshire pigs. Furthermore, histograms of AGE for each subset were left-skewed, indicating that the number of pigs genotyped per year in each subset generally increased from the start of The Maschhoff’s SNP collection platform in 2010 until 2020.

**Table 4.** Descriptive statistics of AGE by subset.

| Subset | Populations | Pigs, n | Mean | SD<sup>1</sup> | Minimum | Maximum |
| --- | --- | --- | --- | --- | --- | --- |
| 1 | Duroc | 16595 | 144.9 | 20.44 | 55 | 171 |
| 2 | Landrace | 15457 | 144.0 | 18.23 | 55 | 171 |
| 3 | Yorkshire | 15772 | 144.2 | 18.46 | 64 | 171 |
| 4 | Crossbred | 8447 | 142.8 | 17.20 | 110 | 164 |
| 5 | Duroc and Landrace | 32066 | 144.5 | 19.41 | 55 | 171 |
| 6 | Duroc and Yorkshire | 32387 | 144.6 | 19.50 | 55 | 171 |
| 7 | Duroc and Crossbred | 25053 | 144.2 | 19.43 | 55 | 171 |
| 8 | Landrace and Yorkshire | 31240 | 144.1 | 18.34 | 55 | 171 |
| 9 | Landrace and Crossbred | 23905 | 143.6 | 17.88 | 55 | 171 |
| 10 | Yorkshire and Crossbred | 24230 | 143.7 | 18.04 | 64 | 171 |
| 11 | Duroc, Landrace and<br>Yorkshire | 47849 | 144.4 | 19.10 | 55 | 171 |
| 12 | Duroc, Landrace, and<br>Crossbred | 40513 | 144.1 | 18.98 | 55 | 171 |
| 13 | Duroc, Yorkshire, and<br>Crossbred | 40837 | 144.2 | 19.06 | 55 | 171 |
| 14 | Landrace, Yorkshire, and<br>Crossbred | 39688 | 143.8 | 18.11 | 55 | 171 |
| 15 | Duroc, Landrace,<br>Yorkshire, and Crossbred | 56296 | 144.2 | 18.84 | 55 | 171 |

<sup>1</sup>SD = standard deviation

**Figure 1.**
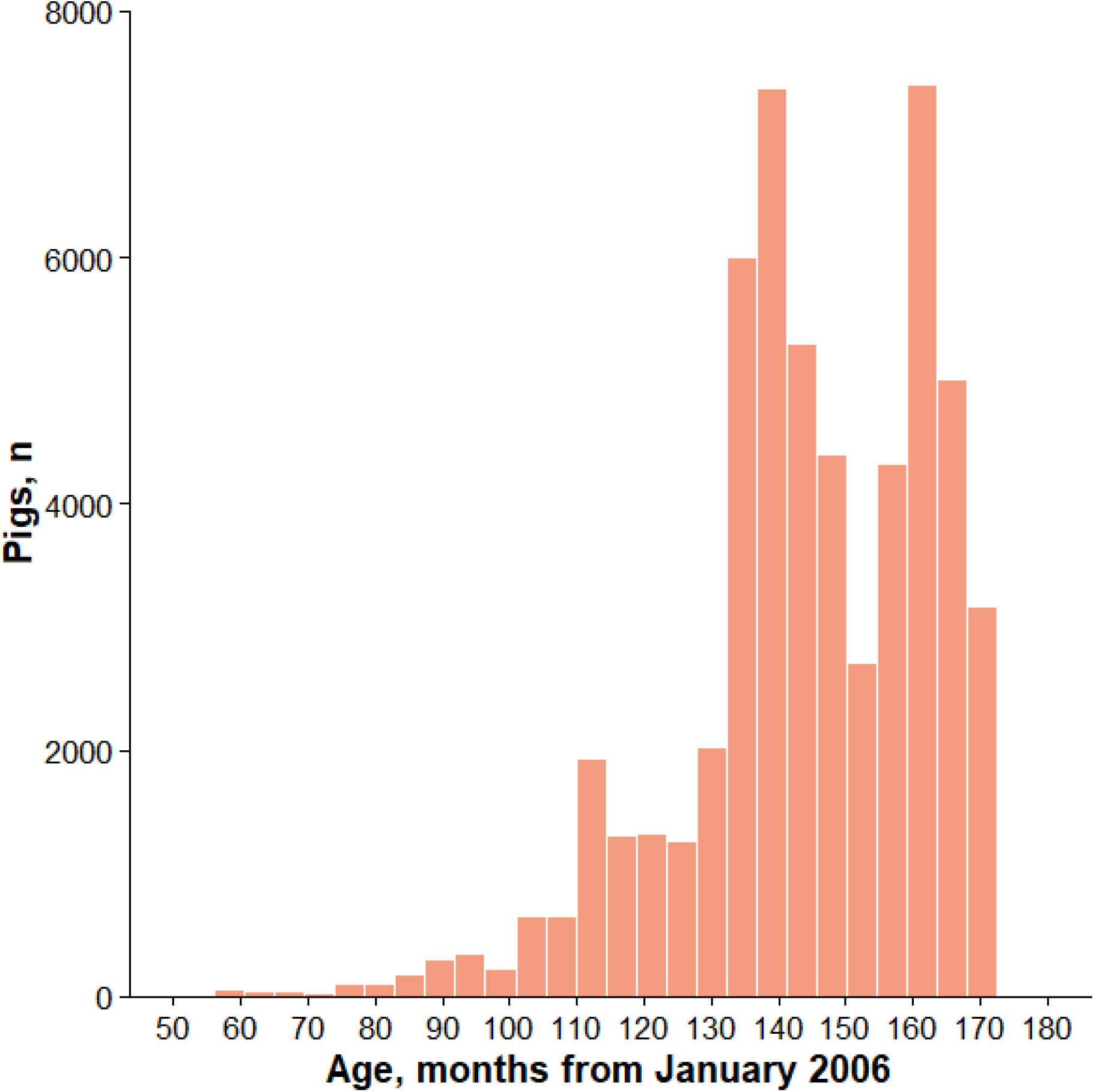
Distribution of AGE for all genotyped pigs. For each pig, AGE was calculated as the number of months between each pig’s birth month and January 2006. A pig with a negative, zero, or positive AGE was born before January 2006, during January 2006, or after January 2006, respectively.

**Figure 2.**
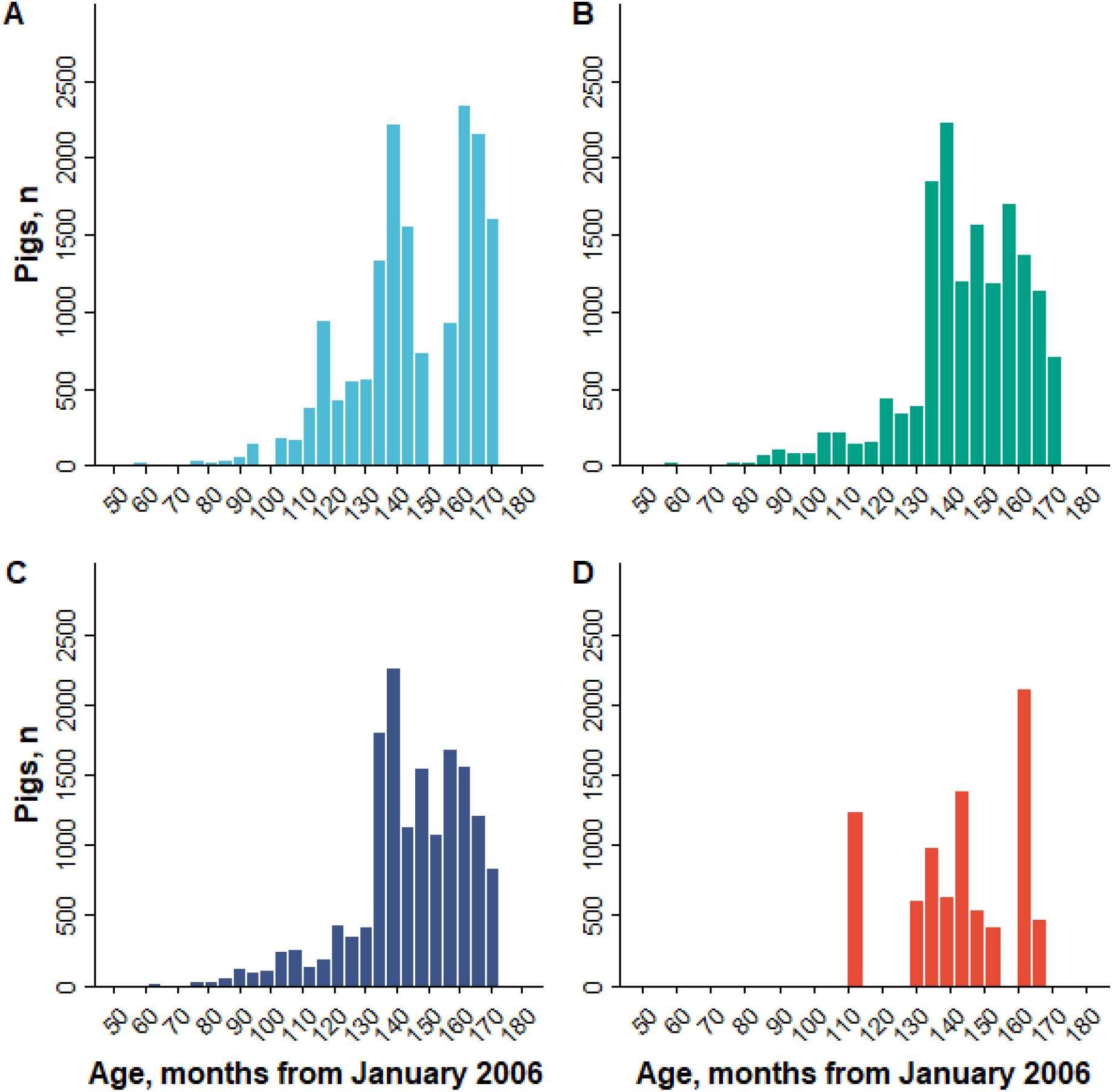
Distributions of AGE for genotyped pigs in Duroc, Landrace, Yorkshire and Crossbreed lines. For each Duroc (**A**), Landrace (**B**), Yorkshire (**C**), and Crossbred (**D**) pig, AGE was calculated as the number of months between each pig’s birth month and January 2006. For example, a pig with an age of 120 was born in January 2016.

Results from the PCA of the GRM containing all pigs from each population (subset 15; Table 3) are presented in Figure 3. By plotting principal component 1 versus principal component 2 for the genomic relatedness of these four populations, four defined clusters were visualized, as expected. In addition, the cluster for the Crossbred population was located halfway between the Duroc population cluster and the Landrace and Yorkshire population clusters along principal component 1 and halfway between the Landrace and Yorkshire population clusters along principal component 2 (Figure 3). McVean et al. [33] postulated that the location of an admixed population of individuals on a PCA plot relative to its source populations directly relates to the admixture proportion of these individuals amongst the source populations. Thus, the location of the Crossbred cluster in Figure 3 confirms approximately a 50/25/25 admixture amongst the Duroc, Landrace, and Yorkshire populations, respectively. This result was to be expected given the design of The Maschhoff’s mating program for their commercial test herd, which mates Duroc sires to Landrace × Yorkshire dams.

**Figure 3.**
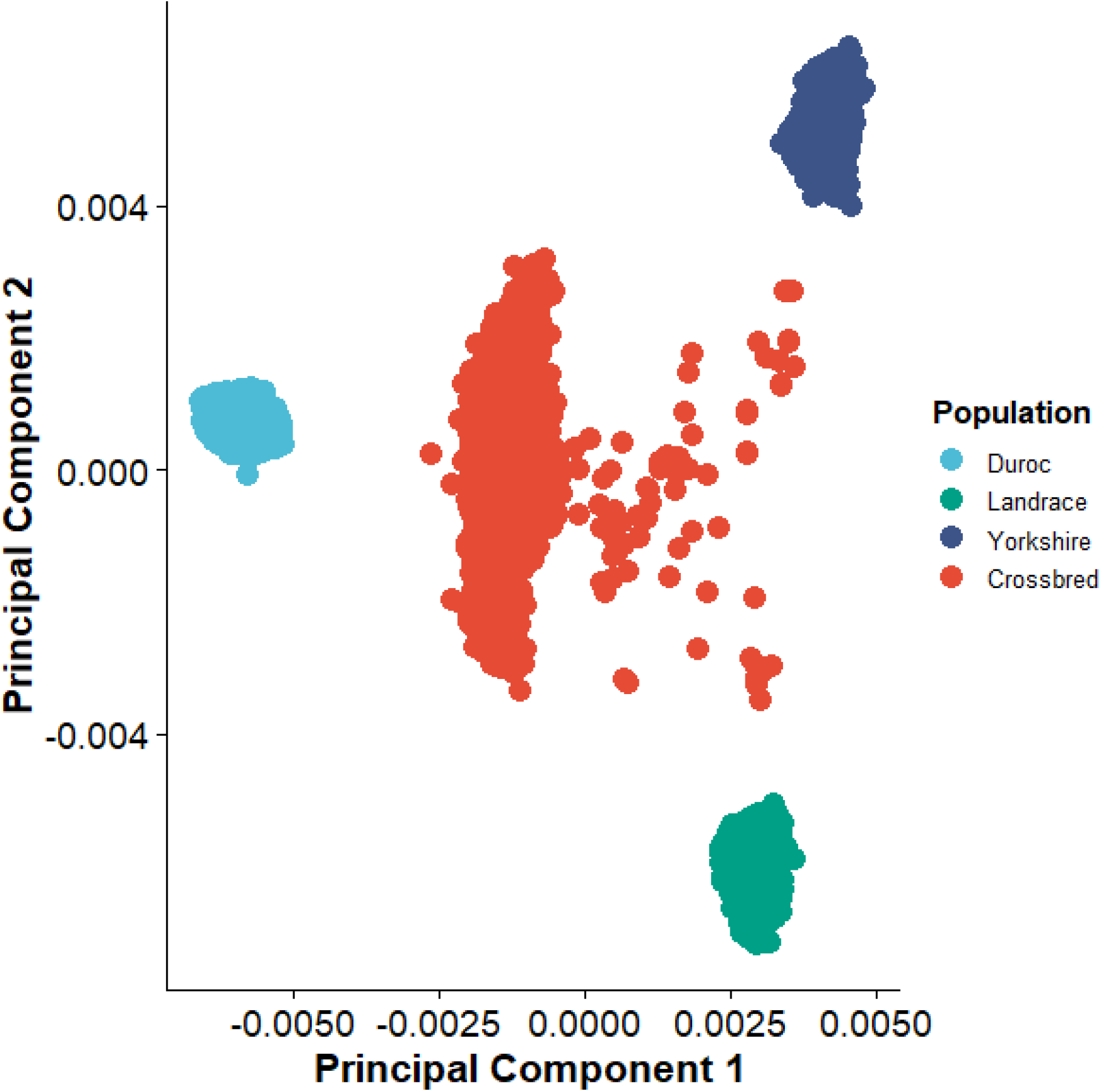
Principal components analysis of GRM containing analyzed genotyped pigs. Four clusters appear in the scatterplot. Individual pigs within each colored cluster constitute a population. Populations of pigs located closer in proximity to one another are genetically more similar.

### Univariate and bivariate variance component estimation

The proportion of variation explained by SNPs of the dependent variable AGE for all 15 subsets is presented in Table 5. These values ranged from 0.81 to 0.94 (Table 5) and were significantly greater than zero (*P* < 0.001) using the likelihood ratio test. Previous GPSM simulations have shown that GPSM PVE is indicative of population demographic history [13]. Given that descriptive statistics and distributions of AGE were generally similar across subsets (Table 4; Figures 1 and 2), these PVE results suggest that the four populations have similar demographics, such as inbreeding, effective population sizes, and pedigree structure. Furthermore, the results of univariate variance component estimation in subsets containing purebred populations (subsets 1, 2, 3, 5, 6, 8, and 11; Table 3) using simulated genotypes are presented in Table 6. Estimated PVEs for subsets with simulated genotypes were generally like those with real data, ranging from 0.83 to 0.93 (Table 6), and were significantly different from zero (*P* < 0.001) using the likelihood ratio test. Between the univariate variance component estimation analyses using real and simulated genotype data, the pedigree structure, AGE values, and number of SNPs were the same for each subset; however, the genomic relationship between each pairwise combination of pigs was different. For the Landrace and Yorkshire populations, the real PVE was only higher than the simulated PVE by 0.02 and 0.01, respectively. However, for the Duroc population, the real PVE was 0.11 higher than the simulated PVE.

**Table 5.**
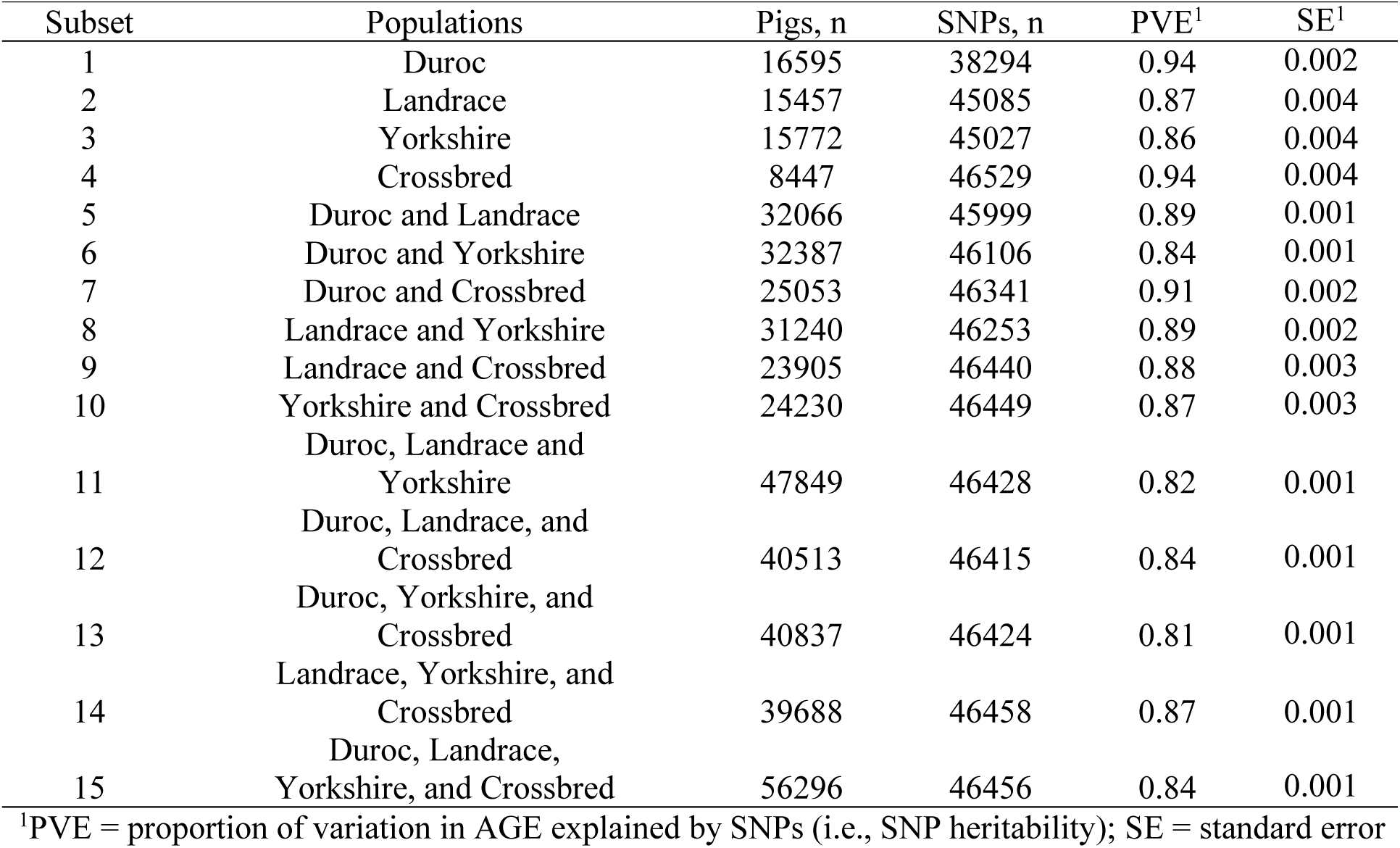
Proportion of variation in AGE explained by SNPs for each subset.

**Table 6.** Proportion of variation in AGE explained by SNPs for each purebred subset using simulated data.

| Subset | Populations | Pigs, n | SNPs, n | PVE<sup>1</sup> | SE<sup>1</sup> |
| --- | --- | --- | --- | --- | --- |
| 1 | Duroc | 16595 | 38286 | 0.83 | 0.005 |
| 2 | Landrace | 15457 | 45090 | 0.85 | 0.004 |
| 3 | Yorkshire | 15772 | 45036 | 0.85 | 0.005 |
| 5 | Duroc and Landrace (Method 1)<sup>2</sup> | 32066 | 46008 | 0.90 | 0.003 |
| 5 | Duroc and Landrace (Method 2) | 32066 | 46008 | 0.88 | 0.003 |
| 6 | Duroc and Yorkshire (Method 1) | 32387 | 46098 | 0.89 | 0.003 |
| 6 | Duroc and Yorkshire (Method 2) | 32387 | 46098 | 0.88 | 0.003 |
| 8 | Landrace and Yorkshire (Method 1) | 31240 | 46260 | 0.91 | 0.002 |
| 8 | Landrace and Yorkshire (Method 2) | 31240 | 46260 | 0.89 | 0.003 |
| 11 | Duroc, Landrace, and Yorkshire (Method 1) | 47849 | 46422 | 0.93 | 0.002 |
| 11 | Duroc, Landrace, and Yorkshire (Method 2) | 47849 | 46422 | 0.90 | 0.002 |

<sup>1</sup>PVE = proportion of variation in AGE explained by SNPs (i.e., SNP heritability); SE = standard error
<sup>2</sup>Method 1 = Genotypes simulated as if populations recently diverged (same founder population); Method 2 = Genotypes simulated as if populations are completely unrelated (different founder populations)

Genetic correlations between AGE for all pairwise combinations of the Duroc, Landrace, Yorkshire, and Crossbred populations (subsets 1 through 4; Table 3) are presented in Table 7. In general, genetic correlations between purebred populations were stronger than genetic correlations between each purebred population and the Crossbred population (Table 7). Each genetic correlation was significantly different from zero (*P* < 0.001) using the likelihood ratio test. Within the purebred subsets (5, 6, and 8; Table 3), the genetic correlation between Landrace and Yorkshire pigs was higher than the genetic correlation between Duroc and Landrace or Duroc and Yorkshire pigs (Table 7). This indicates genomic breeding values for AGE are more similar between Landrace and Yorkshire pigs than between either of the two maternal breeds and the Duroc population. This result was expected, as Landrace and Yorkshire pigs are both breeds known for maternal prolificacy, while Duroc pigs are known for increased efficiency in terminal traits. Amongst each pairwise combination between the Crossbred population and each purebred population (subsets 7, 9, and 10; Table 3), genetic correlations for AGE were highest between the Duroc and Crossbred population and were similar between Crossbred and Landrace or Yorkshire pigs (Table 7). Given that Duroc pigs contribute more genetic material to the Crossbred pigs than the Landrace and Yorkshire pigs, this result was expected.

**Table 7.** Genetic correlations for AGE between each pairwise combination of Populations 1 through 4.

| Subset | Populations | Pigs, n | SNPs, n | $r_G^1$ | SE<sup>1</sup> |
| --- | --- | --- | --- | --- | --- |
| 5 | Duroc and Landrace | 32066 | 45999 | 0.64 | 0.004 |
| 6 | Duroc and Yorkshire | 32387 | 46106 | 0.67 | 0.023 |
| 7 | Duroc and Crossbred | 25053 | 46341 | 0.50 | 0.018 |
| 8 | Landrace and Yorkshire | 31240 | 46253 | 0.80 | 0.017 |
| 9 | Landrace and Crossbred | 23905 | 46440 | 0.38 | 0.021 |
| 10 | Yorkshire and Crossbred | 24230 | 46449 | 0.43 | 0.020 |

<sup>1</sup> $r_G$ = genetic correlation; SE = standard error

Table 8 presents genetic correlations from the bivariate variance component estimation analyses using simulated data. Genetic correlations between each population (subsets 5, 6, and 8; Table 3), using both methods, were not significantly different from zero (*P* > 0.05) using the likelihood ratio test (Table 8). This result suggests that in the absence of artificial selection pressure on economically relevant traits in each population, the correlation for breeding values of AGE is trivial. Moreover, miniscule genetic correlations were observed across both methods used to simulate founder populations; therefore, the length of time from population divergence likely has no effect on genetic correlations in the presence of genetic drift. Thus, the results from this bivariate variance component analysis using simulated founder genotypes strengthen the validity of the assumptions presented above based on real genotype data in genetic lines exposed to artificial selection pressure.

**Table 8.** Genetic correlations for AGE between each pairwise combination of Populations 1 through 3 using simulated genotype data.

| Subgroup | Genetic lines | Pigs, n | SNPs, n | $r_G^1$ | SE<sup>1</sup> |
| --- | --- | --- | --- | --- | --- |
| 5 | Duroc and Landrace (Method 1)<sup>2</sup> | 32066 | 46008 | -0.03 | 0.028 |
| 5 | Duroc and Landrace (Method 2) | 32066 | 46008 | -0.02 | 0.058 |
| 6 | Duroc and Yorkshire (Method 1) | 32387 | 46098 | 0.06 | 0.029 |
| 6 | Duroc and Yorkshire (Method 2) | 32387 | 46098 | 0.03 | 0.057 |
| 8 | Landrace and Yorkshire (Method 1) | 31240 | 46260 | -0.02 | 0.028 |
| 8 | Landrace and Yorkshire (Method 2) | 31240 | 46260 | -0.01 | 0.056 |

<sup>1</sup> $r_G$ = genetic correlation; SE = standard error
<sup>2</sup>Method 1 = Genotypes simulated as if populations recently diverged (same founder population); Method 2 = Genotypes simulated as if populations diverged several years ago (different founder populations)

### Detecting polygenic selection with Generation Proxy Selection Mapping (GPSM)

The number of significant SNPs (*Q* < 0.10) associated with AGE for each subset is presented in Table 9. Although the distribution of AGE for each subset was left-skewed and non-normal (Table 4; Figures 1 and 2), the GPSM *P*-values for independent SNP genotype association tests with AGE were well calibrated (Figure 4). For example, *P*-values for null SNPs, which were deemed non-significant by GPSM, closely followed the expected uniform distribution, while SNPs that were significantly associated with AGE deviated from this expectation (Figure 4). This result suggests that departures from normality in the dependent variable (i.e., generation-proxy) in a GPSM analysis does not produce spurious associations between AGE and genotype. Generation proxy selection mapping identified 49 to 854 significant SNPs (Table 9) depending on subset. The number of significant associations generally increased as the number of samples in the subset increased, as expected, due to increased power of the genome wide association analysis.

**Table 9.** Number of SNPs significantly associated with AGE for each subset.

| Subset | Populations | Pigs, n | SNPs, n | Significant SNPs, n<sup>1</sup> |
| --- | --- | --- | --- | --- |
| 1 | Duroc | 16595 | 38294 | 100 |
| 2 | Landrace | 15457 | 45085 | 147 |
| 3 | Yorkshire | 15772 | 45027 | 138 |
| 4 | Crossbred | 8447 | 46529 | 49 |
| 5 | Duroc and Landrace | 32066 | 45999 | 371 |
| 6 | Duroc and Yorkshire | 32387 | 46106 | 527 |
| 7 | Duroc and Crossbred | 25053 | 46341 | 148 |
| 8 | Landrace and Yorkshire | 31240 | 46253 | 177 |
| 9 | Landrace and Crossbred | 23905 | 46440 | 172 |
| 10 | Yorkshire and Crossbred | 24230 | 46449 | 182 |
| 11 | Duroc, Landrace and<br>Yorkshire | 47849 | 46428 | 702 |
| 12 | Duroc, Landrace, and<br>Crossbred | 40513 | 46415 | 533 |
| 13 | Duroc, Yorkshire, and<br>Crossbred | 40837 | 46424 | 609 |
| 14 | Landrace, Yorkshire, and<br>Crossbred | 39688 | 46458 | 274 |
| 15 | Duroc, Landrace,<br>Yorkshire, and Crossbred | 56296 | 46456 | 854 |

<sup>1</sup> $Q < 0.10$

**Figure 4.**
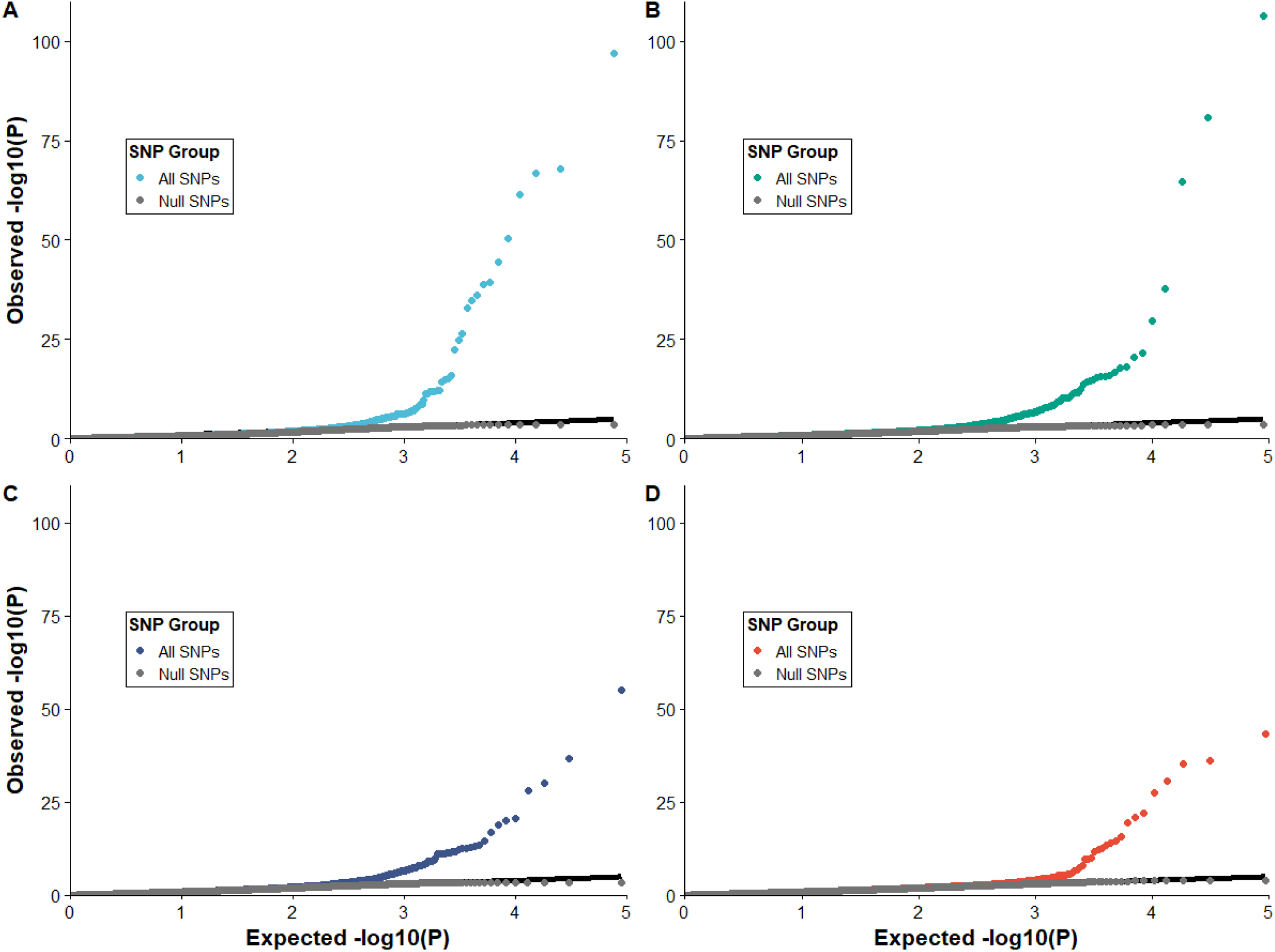
Q-Q plots for GPSM P-values from genome-wide association analyses of SNP genotype on AGE. Null SNPs (non-significant) closely followed a uniform distribution, while GPSM significant SNPs deviated from the expected uniform distribution for Duroc (**A**), Landrace (**B**), Yorkshire (**C**), and Crossbred (**D**).

There were 100, 147, 138, and 49 significant SNPs identified by GPSM representing 0.26, 0.33, 0.31, and 0.11% of the total number of autosomal SNPs for the Duroc, Landrace, Yorkshire, and Crossbred populations, respectively (subsets 1 through 4; Table 9). However, when all purebred pigs were combined into a single subset (subset 11; Table 3), GPSM identified 702 significant associations (1.51% of autosomal loci; Table 9). Moreover, the addition of the Crossbred pigs to subset 11, which created subset 15 (Table 3), allowed GPSM to identify 854 significant associations (1.84% of autosomal loci; Table 9). As mentioned above, the efficacy of GPSM analyses depends on the power of the genome-wide association analyses. Thus, as more samples of SNP genotype information on a particular population of pigs are accumulated, more SNP genotypes that are associated with AGE can be detected using the GPSM method.

Manhattan plots of −*log*_10_(*Q*) values for the associations between SNP genotypes and AGE in the Duroc, Landrace, Yorkshire, and Crossbred populations (subsets 1 through 4; Table 3) are presented in Figure 5. For each population, a plot is presented with a full (Figure 5A-5D) and a truncated Y-axis, which ranged in −*log*_10_(*Q*) values from 0 to 10 (Figure 5E-5H). Within each subset, several significant associations between SNP genotype and AGE were identified on each chromosome by GPSM (Figure 5). When viewing the Manhattan plots for each genetic line with truncated Y-axes, genome-wide nature of the significant associations becomes more pronounced (Figure 5E-5H).

**Figure 5.**
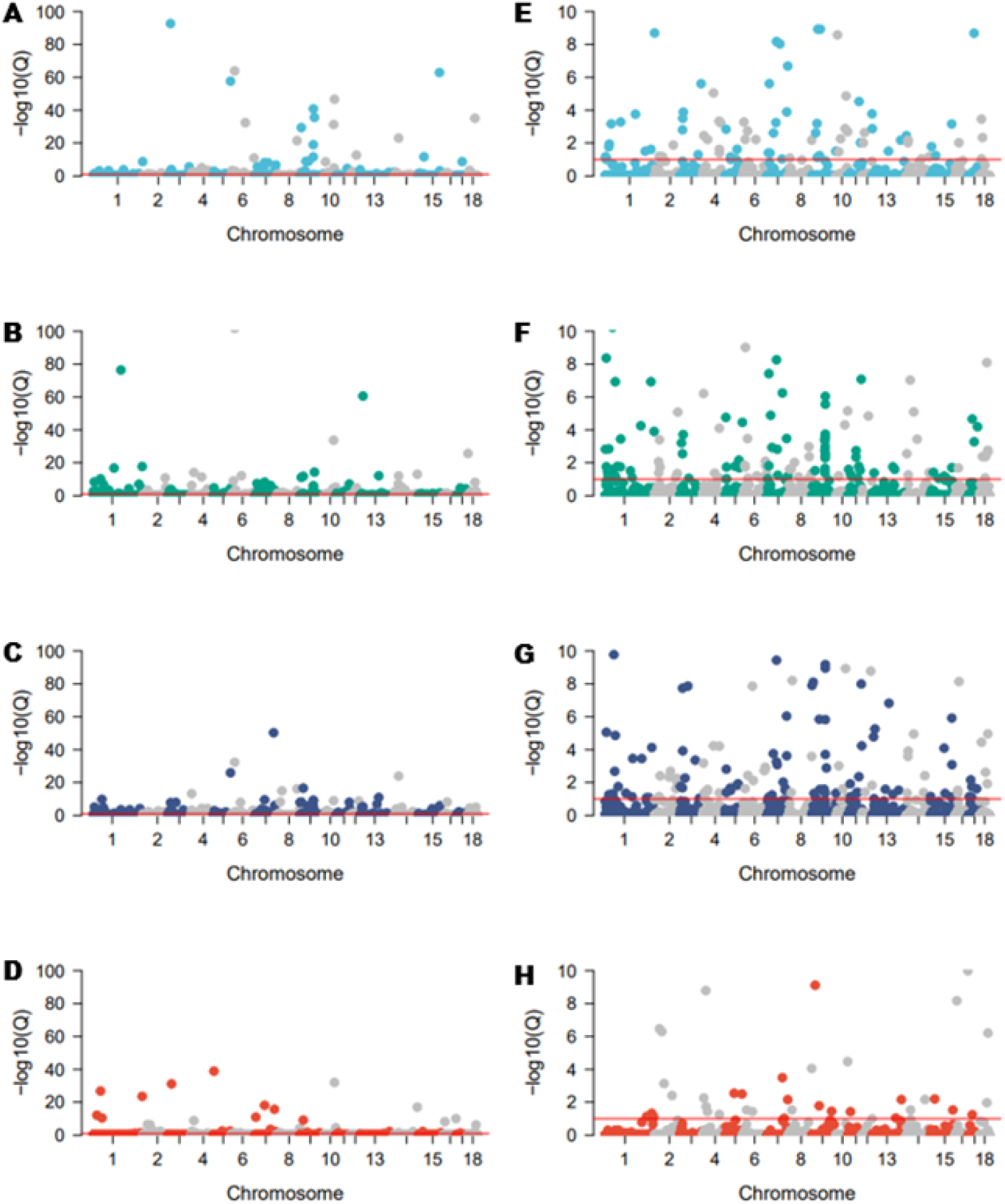
Manhattan plots of GPSM Q-values for the association between SNP genotype and AGE. Significant GPSM SNPs were found on each chromosome, and -log_10_(*Q*-values) and are shown in Manhattan plots with full Y-axes for Duroc (**A**), Landrace (**B**), Yorkshire (**C**), and Crossbred (**D**). Truncated Y-axes from 0 to 10 -log_10_(*Q*-values) reveal polygenic nature of selection in Duroc (**E**), Landrace (**F**), Yorkshire (**G**), and Crossbred (**H**).

Distributions of allele substitution effects are plotted in Figure 6. For each population, the distributions of allele substitution effects for significant GPSM SNPs followed a bimodal distribution, with central values for each of the peaks located below and above zero (Figure 6). In contrast, distributions of allele substitution effects for null (non-significant) GPSM SNPs followed a normal distribution, with mean values located near zero (Figure 6). Allele substitution effects for SNPs in each population that were significantly different than zero were converted to absolute values to interpret differences in magnitude of this parameter across populations. Duroc pigs had the highest mean absolute value of age allele substitution effects for significant SNPs (2.70 months) and mean absolute values of allele substitution effects in significant SNPs were similar between the Landrace (1.66 months), Yorkshire (1.55 months), and Crossbred (1.80 months) populations. The range in absolute values of allele substitution effects in GPSM significant SNPs was considerable, depending on population. For example, in the Duroc and Crossbred populations, these ranges were 1.00 to 13.32 months and 0.70 to 14.39 months, respectively. However, for the Landrace and Yorkshire populations, these ranges were smaller (0.71 to 6.64 and 0.71 to 6.00 months, respectively) but were similar between the two populations.

**Figure 6.**
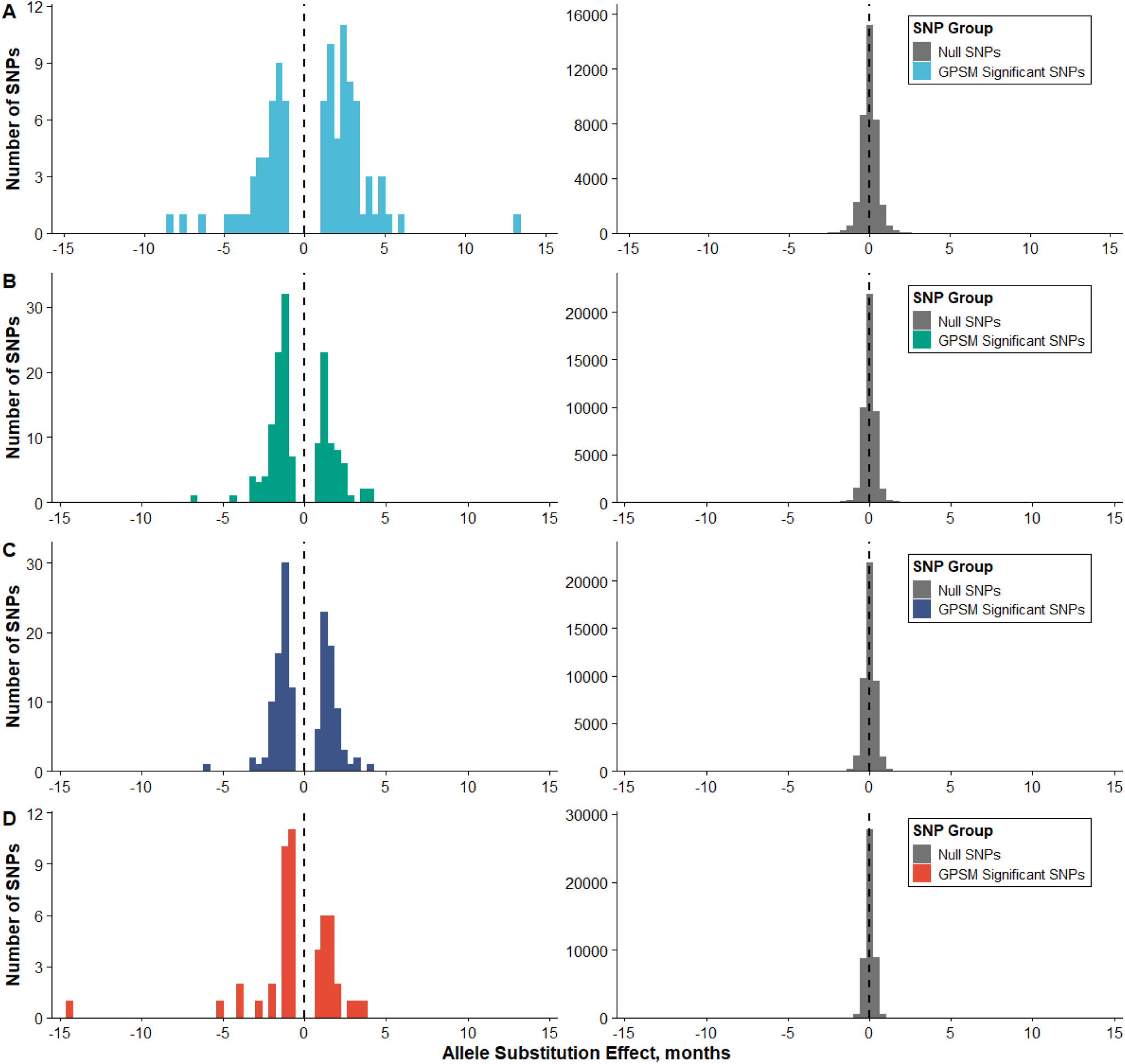
Distribution of allele substitution effects for null and GPSM significant SNPs. For Duroc (**A**), Landrace (**B**), Yorkshire (**C**), and Crossbred (**D**), null SNPs (non-significant) were normally distributed with a mean near zero, while GPSM significant SNPs followed a bimodal distribution with central values for each peak located above and below zero.

Results from GPSM analyses using randomly simulated founder genotypes are presented in Table 10. Out of the 11 GPSM runs on the simulated data, GPSM falsely identified significant associations with AGE in seven analyses (Table 10). However, in these analyses, a very small number of spurious associations were detected (Table 10), corresponding to a range of error rates between 0.0022 to 0.0152% (Table 10), which are negligible. Analyses that used Method 2 to simulate founder genotypes, which simulated completely different founder genotypes for each population, had a higher number of spurious associations than Method 1, which simulated a single founder population for all three purebred populations. In commercial pig populations, however, divergence likely happened in a scenario that resembles a blending of Methods 1 and 2; thus, the higher error rate in Method 2 could be inflated compared to reality in the swine industry. Nevertheless, these results suggest that GPSM is robust to allele frequency changes due to genetic drift over time.

**Table 10.**
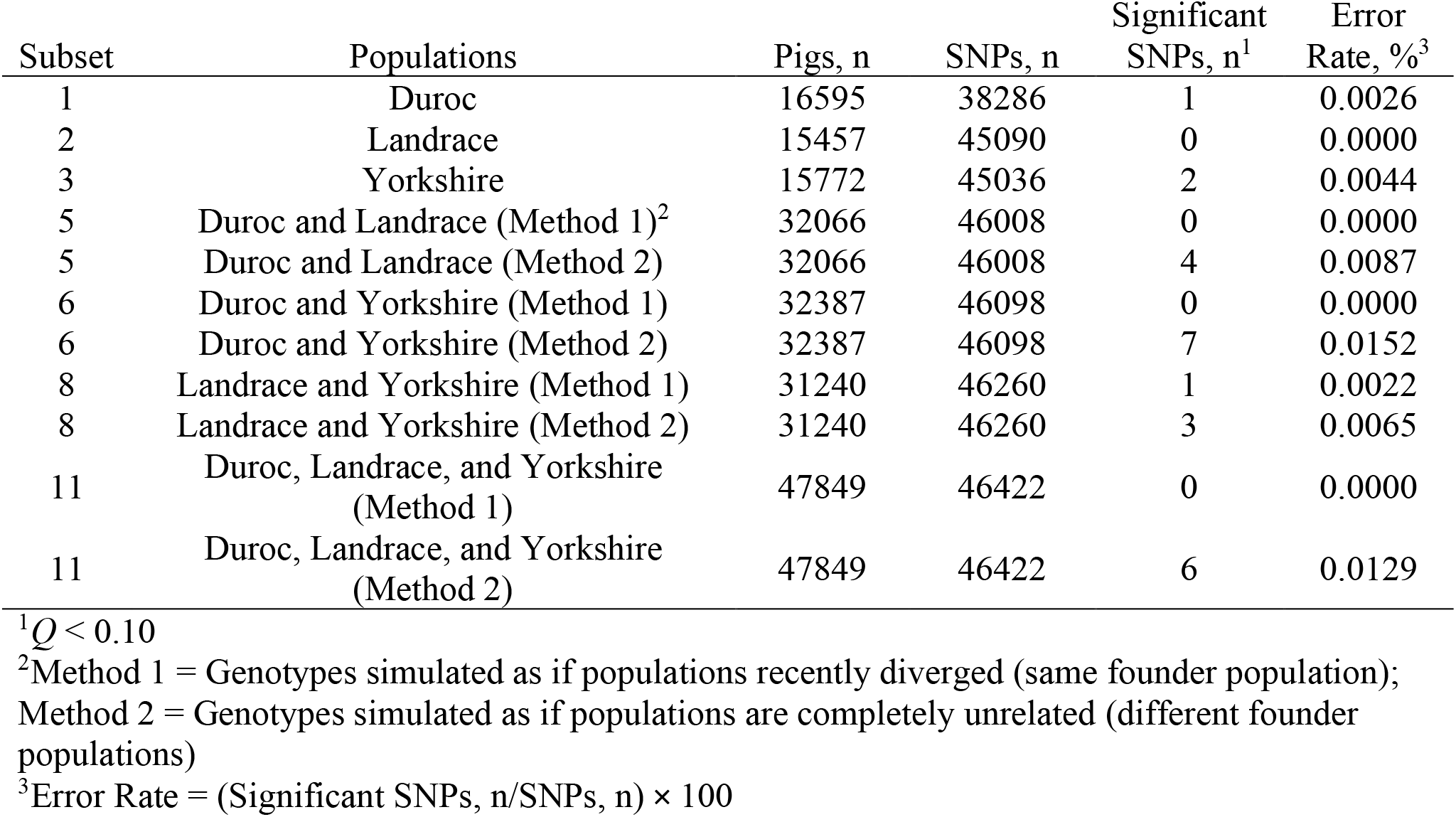
Number of SNPs significantly associated with AGE for each subset using randomly simulated genotype data.

| Subset | Populations | Pigs, n | SNPs, n | Significant SNPs, n<sup>1</sup> | Error Rate, %<sup>3</sup> |
| --- | --- | --- | --- | --- | --- |
| 1 | Duroc | 16595 | 38286 | 1 | 0.0026 |
| 2 | Landrace | 15457 | 45090 | 0 | 0.0000 |
| 3 | Yorkshire | 15772 | 45036 | 2 | 0.0044 |
| 5 | Duroc and Landrace (Method 1)<sup>2</sup> | 32066 | 46008 | 0 | 0.0000 |
| 5 | Duroc and Landrace (Method 2) | 32066 | 46008 | 4 | 0.0087 |
| 6 | Duroc and Yorkshire (Method 1) | 32387 | 46098 | 0 | 0.0000 |
| 6 | Duroc and Yorkshire (Method 2) | 32387 | 46098 | 7 | 0.0152 |
| 8 | Landrace and Yorkshire (Method 1) | 31240 | 46260 | 1 | 0.0022 |
| 8 | Landrace and Yorkshire (Method 2) | 31240 | 46260 | 3 | 0.0065 |
| 11 | Duroc, Landrace, and Yorkshire (Method 1) | 47849 | 46422 | 0 | 0.0000 |
| 11 | Duroc, Landrace, and Yorkshire (Method 2) | 47849 | 46422 | 6 | 0.0129 |

<sup>1</sup> $Q < 0.10$
<sup>2</sup>Method 1 = Genotypes simulated as if populations recently diverged (same founder population); Method 2 = Genotypes simulated as if populations are completely unrelated (different founder populations)
<sup>3</sup>Error Rate = (Significant SNPs, n/SNPs, n) × 100

The number of shared significant GPSM SNPs across subsets 1 through 4 is presented in Figure 7. Forty-two, 22, and 4 SNPs significantly associated with AGE were shared across at least two, three, or four populations, respectively (Figure 7). Twenty-five GPSM SNPs were unique to the Landrace and Yorkshire populations, which was considerably larger than the number of unique GPSM SNPs identified between all other pairwise combinations of each individual population (Figure 7). In addition, 13 GPSM SNPs were unique to the purebred populations. However, only 2 to 4 GPSM SNPs were unique to subsets of three populations where the Crossbred pigs were included (subsets 12 through 14; Figure 7). Large allele substitution effect SNPs that were significantly associated with AGE in the Duroc, Landrace, Yorkshire, and Crossbred populations are shown in Table 11. In general, most of the significant SNPs with the 10 largest absolute values for allele substitution effects were significant in at least one other subset (Table 11). In the Crossbred population, only 2 of the top 10 large effect SNPs were unique to Crossbred pigs (3, 4, and 1 out of 10 were significant across at least 2, 3 and 4 subsets, respectively; Table 11). In addition, certain SNPs exhibited large allele substitution effects across multiple subsets. For example, GPSM estimated an allele substitution effect for SNP 39502 of 6.19, -6.64, and 4.03 months (Table 11) in the Duroc, Landrace, and Yorkshire populations, respectively, which were 11.9, 18.4, and 10.9 SD above, below, and above the mean allele substitution effect within each subset, respectively.

**Figure 7.**
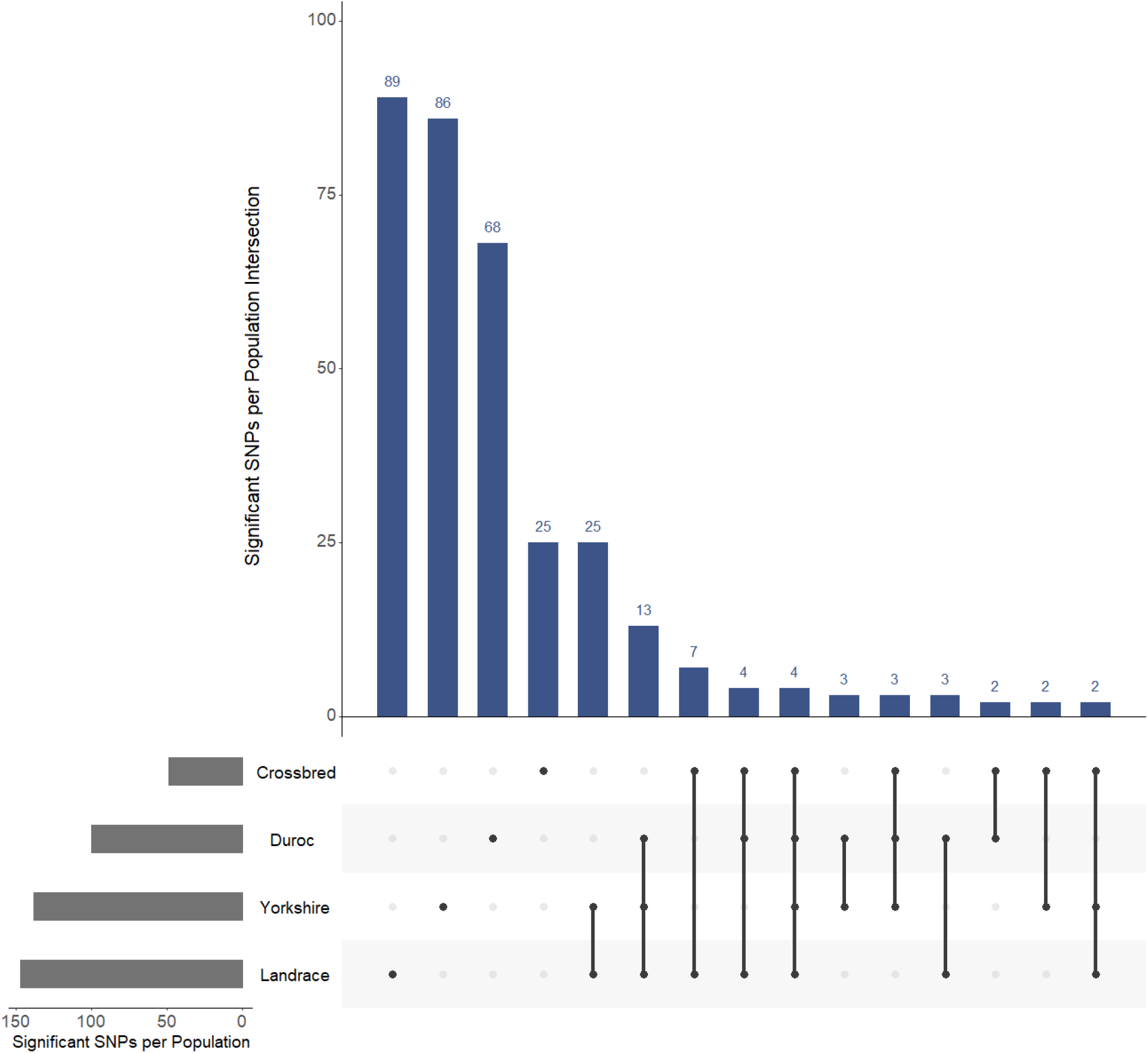
UpSet plot depicting the number of GPSM significant SNPs across populations. Each vertical blue bar shows the number of GPSM significant SNPs unique to a single population (25 to 89 SNPs), unique across two populations (2 to 25 SNPs), unique across three populations (2 to 13 SNPs), or unique across all four populations (4 SNPs). Horizontal gray bars present the number of GPSM significant SNPs in each genetic line.

**Table 11.** Ten SNPs significantly associated with AGE with the largest absolute values for allele substitution effect within subsets 1 through 4.^1^

| Subset | Populations | SNP Identifier | MAF | ASE | SE | Q-value | Number of subsets where significant |
| --- | --- | --- | --- | --- | --- | --- | --- |
| 1 | Duroc | 3420 | 0.24 | 13.32 | 0.310 | 1.49E-102 | 3 |
|  |  | 41017 | 0.02 | 8.21 | 0.648 | 3.54E-33 | 1 |
|  |  | 30819 | 0.07 | 7.71 | 0.368 | 2.05E-93 | 2 |
|  |  | 35689 | 0.09 | 6.49 | 0.390 | 2.34E-58 | 3 |
|  |  | 39502 | 0.47 | 6.19 | 0.353 | 1.24E-64 | 3 |
|  |  | 18513 | 0.02 | 5.16 | 0.643 | 2.28E-12 | 1 |
|  |  | 30855 | 0.13 | 4.87 | 1.025 | 1.56E-03 | 1 |
|  |  | 49794 | 0.38 | 4.79 | 0.340 | 1.56E-41 | 3 |
|  |  | 31005 | 0.11 | 4.66 | 0.880 | 1.31E-04 | 1 |
|  |  | 6055 | 0.08 | 4.60 | 0.370 | 5.40E-32 | 3 |
| 2 | Landrace | 39502 | 0.30 | 6.64 | 0.302 | 1.76E-102 | 3 |
|  |  | 14465 | 0.02 | 4.40 | 0.518 | 9.00E-14 | 3 |
|  |  | 1063 | 0.46 | 3.95 | 0.207 | 3.69E-77 | 1 |
|  |  | 22747 | 0.31 | 3.81 | 0.333 | 2.43E-26 | 1 |
|  |  | 10745 | 0.42 | 3.52 | 0.206 | 2.66E-61 | 2 |
|  |  | 38925 | 0.04 | 3.51 | 0.840 | 1.55E-02 | 1 |
|  |  | 6055 | 0.18 | 3.08 | 0.238 | 2.03E-34 | 3 |
|  |  | 485 | 0.23 | 3.08 | 0.325 | 1.71E-17 | 2 |
|  |  | 39264 | 0.28 | 3.03 | 0.418 | 9.58E-10 | 1 |
|  |  | 8174 | 0.29 | 3.00 | 0.456 | 8.35E-08 | 2 |
| 3 | Yorkshire | 45804 | 0.02 | 6.00 | 0.664 | 1.14E-15 | 1 |
|  |  | 39502 | 0.36 | 4.03 | 0.316 | 4.80E-33 | 3 |
|  |  | 19756 | 0.17 | 3.40 | 0.486 | 7.20E-09 | 1 |
|  |  | 35689 | 0.28 | 3.28 | 0.284 | 1.24E-26 | 3 |
|  |  | 9883 | 0.39 | 3.05 | 0.385 | 1.20E-11 | 2 |
|  |  | 8174 | 0.14 | 3.03 | 0.437 | 1.02E-08 | 2 |
|  |  | 41553 | 0.43 | 2.96 | 0.188 | 5.05E-51 | 1 |
|  |  | 30156 | 0.04 | 2.87 | 0.790 | 9.16E-02 | 1 |
|  |  | 34751 | 0.05 | 2.52 | 0.461 | 5.84E-05 | 3 |
|  |  | 34530 | 0.18 | 2.38 | 0.277 | 6.05E-14 | 2 |
| 4 | Crossbred | 30819 | 0.07 | 14.39 | 0.372 | 1.81E-102 | 2 |
|  |  | 14465 | 0.02 | 5.10 | 0.536 | 9.90E-18 | 3 |
|  |  | 3834 | 0.04 | 4.01 | 0.344 | 1.58E-27 | 2 |
|  |  | 36398 | 0.18 | 4.00 | 0.288 | 1.25E-39 | 2 |
|  |  | 31018 | 0.48 | 3.52 | 0.281 | 7.71E-32 | 1 |
|  |  | 6187 | 0.40 | 3.06 | 0.241 | 1.00E-32 | 3 |
|  |  | 3420 | 0.43 | 2.99 | 0.271 | 2.72E-24 | 3 |
|  |  | 43184 | 0.08 | 2.77 | 0.282 | 7.68E-19 | 4 |
|  |  | 33882 | 0.07 | 2.09 | 0.290 | 1.63E-09 | 3 |
|  |  | 4012 | 0.13 | 2.00 | 0.258 | 4.15E-11 | 1 |

<sup>1</sup>MAF = minor allele frequency; ASE = allele substitution effect; SE = standard error

Positional candidate genes located within 100 kb upstream or downstream of significant SNPs of the chromosome 9 association found in all three purebred populations revealed four protein-coding genes, *MIOS, RPA3, UMAD1*, and *GLCCI1*.

## Discussion

Polygenic selection on quantitative traits, induces small changes in allele frequencies at numerous loci across the genome over time [13,34,35]. Detection of this polygenic selection due to artificial selection over time for traits with complex architectures was the focus of this study. The increasing abundance of genomic information from SNP arrays [36] has allowed many researchers to study changes in genotypic and allelic frequency in commercial and indigenous global pig populations [2,3,15,37,38]. The rapid increase in studies in this area has given rise to new analytical methods to detect large and small selection signatures across in the genome of commercially-reared livestock species, such as Generation Proxy Selection Mapping (GPSM) [13,14]. In the present study, GPSM was used to estimate variance components and SNP genotype associations with the dependent variable AGE, which was calculated as the difference in months from January 2006 in a large commercial population of pigs that was comprised of three distinct pure populations (Duroc, Landrace, and Yorkshire) and a crossbred population comprised of the three pure populations. We found that the genomic relationship matrix accounted for confounding due to pedigree and population structure consistently across seven gene drop simulations, as false positive rates ranged between 0% to 0.015% (Table 10).

The proportion of variation in age explained by the GRM ranged from 0.81 to 0.94. Rowan et al. [13] stated, based off of simulations of genotypes from random mating versus selection analyzed with GPSM, that PVE is a function of the number of generations of selection, the number of total crosses per generation, and the genotype sampling scheme (even or uneven across generations). Our results are consistent with these conclusions as similar results across univariate variance component analyses using real and simulated genotype data indicated pedigree structure and the distribution of AGE were the main determinants of PVE (Tables 5 and 6). The difference between the gene-drop simulation PVE and observed PVE was small for the Landrace and Yorkshire population but was 0.11 for the Duroc population. The main difference between simulations and observed data was the presence of selection, suggesting that the Duroc population was under stronger selection compared to the Landrace and Yorkshire populations. Selection indices for Duroc terminal populations generally consist of only traits related to growth, carcass and feed consumption, while selection indices for maternal lines consist of the previously stated traits and additional traits related to maternal prolificacy. Thus, overall genetic merit likely improved at a slower pace in the maternal populations compared to the Duroc population as a result of the added traits in the selection index. Further, Rowan et al. [13] found smaller PVEs across three cattle populations [PVE = 0.52, 0.59, and 0.46 in Red Angus (n = 15,295), Simmental (n = 15,350), and Gelbvieh (n = 13,031) populations, respectively] of similar sample sizes to the purebred populations in the current study. Differences between cattle and pigs in overall structure of the genetic selection programs related to the above factors likely contributed to the large difference between the PVEs reported by Rowan et al. [13] and the present study.

The estimation of genetic correlations between each pairwise combination of the Duroc, Landrace, Yorkshire, and Crossbred populations confirmed our assumptions of similarity (or dissimilarity) between populations in the genomic influence of AGE. While genetic correlations between AGE of two populations can be hard to interpret from a biological standpoint, a genetic correlation near 1 suggests a high proportion of autosomal loci that are statistically associated with AGE undergoing similar changes in allelic frequency over time, with a genetic correlation between 0 and -1 suggesting the contrary (dissimilar or antagonistic changes in allele frequency in SNPs associated with AGE over time). The results of the simulation analysis, where randomly generated founder pig SNP genotypes are randomly dropped through the real pedigree of each population (mimicking genetic drift), validate this assumption, as genetic correlations between populations were not significantly different from zero (regardless of the most recent common ancestor in simulations) using the likelihood ratio test (*P* > 0.05; Table 8). Selection objectives within The Maschhoffs are highly similar between the Landrace and Yorkshire populations and are the most dissimilar between the Duroc and each of the Landrace and Yorkshire populations. Estimated genetic correlations in the present study followed this premise, as the estimated genetic correlations between the two maternal breeds was higher than those estimated between the Duroc population and either the Landrace or Yorkshire populations (Table 7). However, across all four populations, the genetic correlations were significantly larger than zero, indicating that selected loci are similar across populations. This is supported by GPSM associations, as most strong associations were identified in multiple populations (Figure 7) and there was a general increase in associations when pooling populations (subsets 5 through 15; Table 9).

Generation Proxy Selection Mapping analyses in the current study identified hundreds of SNPs significantly associated with AGE (*Q* < 0.10) in most populations (Table 9). There was a wide range in the number of pigs in each subset used in the GPSM analyses (Table 9). The GPSM method, as stated above and in other studies [13,14], is a genome-wide association analysis, which are more powerful in the detection of SNP genotypes associated with a particular phenotype as the number of samples in the population increases, due to increased precision in estimating allele substitution effects at a particular SNP [36]. This inherent attribute of GWAAs contributed to the vast differences in the number of significant associations between SNP genotypes and AGE across subsets, as the number of significant SNPs showed a general increase with sample size (Table 9). However, this is only the case if the same loci are increasing in frequency across the different populations. The overwhelming majority of autosomal SNPs for each subgroup were not associated with AGE, according to GPSM results (98.2 to 99.9% of the autosomal loci; Table 9). However, GPSM detected several SNPs significantly associated with AGE on each chromosome (Figure 5). In addition, the nature of the genome-wide associations with AGE indicates that selection in these populations is likely polygenic (Figure 5E-5F).

A number of SNPs were detected by GPSM across at least two populations (Figure 7). Upon visual assessment of Manhattan plots of GPSM *Q*-values for each population, several regions along the autosomal genome that expressed similar patterns of GPSM significance across populations were identified (Figure 5). Of particular interest, from the chromosome 9 region associated with selection in all three purebred populations, the four candidate genes (*MIOS, RPA3, UMAD1*, and *GLCCI1*) are all differentially expressed in ovarian tissues [39]. Most notably, *MIOS*, which is commonly referred to as the “missing oocyte gene”, is well known for its role in meiosis regulation of oocyte development. In a study using *Drosophila*, a mutation in the *MIOS* gene caused erroneous oocyte development. Instead of stimulating progression through each stage of meiosis, the described mutation caused oocyte progression towards polyploid nurse cells as opposed to fully functional, mature haploid gametes [40]. While there are no known studies that have evaluated the impact of mutations in the “missing oocyte” gene in pigs, results from the present study suggest selection pressure in The Maschhoff’s genetic program has had a significant effect on regions of the pig genome that influence fertility. As a litter-bearing species, pig breeders routinely place selection pressure on litter traits such as total born and number born alive, especially in Landrace and Yorkshire pig populations. In addition, not only does selecting young replacement animals influence allele frequencies at quantitative trait loci, but decisions on which animals to cull likely have similar effects. For example, gilts or sows in breeding populations that fail to express estrus cyclicity, conceive or farrow litters, or return to estrus within a reasonable period post-weaning are typically removed from the herd. Selection or culling of breeding animals due to reproductive performance and fertility issues, respectively, likely caused allele frequency changes at loci near these 4 genes on chromosome 9. However, further quantitative trait association studies and bioinformatics analyses are required to confirm this assumption. Nonetheless, the detection of significant associations across the autosomal genome in each of the Duroc, Landrace, Yorkshire, and Crossbred populations indicates artificial selection is influencing numerous genes in each of these populations of pigs. Further, to see an increase of power when pooling data across populations and shared signal across populations, there must be common causal variants (or at a minimum, casual genes) segregating in the populations, and the variants must be responding to similar selection objectives. Thus, concordant traits across selection indices for maternal and terminal pig breeds are likely influenced by the same quantitative trait loci in the genome of each breed.

We confirmed GPSM is robust in separating changes in allele frequency due to genetic drift and artificial selection, through simulations. In each of the 11 gene-drop simulations, GPSM found very few spurious associations between SNP genotype and AGE (Table 10). Rowan et al. [13] identified false positives as significant at a rate of one SNP per 100,000 tests, which was similar to the current study.

Except for two outliers [SNP 30819 in the Crossbred population (14.39 months) and SNP 3420 in the Duroc population (13.32 months); Table 11], the absolute values of allele substitution effects ranged between 0 and 8.21 months. Mean absolute values for AGE allele substitution effects in significant SNPs were higher in the Duroc population (2.70 months) than the other two purebred populations (1.66 and 1.55 months for the Landrace and Yorkshire populations, respectively). This suggests that selection intensity is greater in the Duroc population, which induces larger changes in allele frequency over shorter periods of time than in the maternal breed populations. Selection in the Duroc population within The Maschhoff’s has been focused on traits that increase the efficiency of terminal commercial progeny, such as increased growth and feed efficiency, decreased backfat depth, and increased carcass lean content. Growth and carcass traits, in general, have more accurate genetic predictions due to their moderate to large heritabilities, which increases selection response as opposed to selection on maternal traits such as number of piglets born alive and litter weaning weight (traits that are emphasized in The Maschhoff’s Landrace and Yorkshire populations). Moreover, as stated previously, the maternal selection indices consisted of more traits, which could have decreased the rate of genetic progress for any single trait relative to an index consisting of fewer traits. This difference in breeding objective between these two groups of genetic lines is likely responsible for higher AGE allele substitution effects in the Duroc pigs. Crossbred pigs in The Maschhoff’s genetic selection program are not exposed to direct selection pressure. Rather, artificial selection occurs in the three genetic lines that constitute the genetic makeup of the Crossbred population. Mean absolute values for AGE allele substitution effects in significant SNPs in the Crossbred pigs (1.80 months) were similar to values reported for the three pure populations, suggesting that selection in the three purebred populations also changes allele frequencies in the Crossbred population at similar rates. However, it must be noted that genotype samples in Crossbred population were collected over a span of approximately 4 years, as opposed to a span of approximately 10 years in the three purebred populations (Table 2).

## Conclusions

We evaluated Generation Proxy Selection Mapping as an analytical method for detecting large and small signatures of artificial selection in a large commercial population of pigs from three purebred populations and one crossbred population. Numerous significant SNPs were detected across the genome in each genetic line, indicating that GPSM is effective in detecting changes in pig genomes due to polygenic selection over relatively short time scales (~4 to 10 years). In addition, simulations proved that GPSM is well-calibrated to distinguish between allele frequency changes over time resulting from genetic drift or artificial selection. Several SNPs were identified as significantly associated with AGE across multiple populations, indicating similar selection objectives, genetic architectures, and causal variants underlying quantitative traits influencing allele frequencies at loci in each population over time. Results from this analysis and future analyses using GPSM could give valuable insight into biological mechanisms underlying and responding to selection on quantitative phenotypes in the commercial swine industry. Lastly, SNPs identified as significantly associated with AGE have the potential to serve as indicators of genomic regions to highlight in the development of genetic prediction models and selection schemes in swine breeding programs.

## Declarations

### Ethics approval and consent to participate

Because phenotypic records and tissue samples were collected as part of routine livestock production practices, and were obtained from an existing industry database, ACUC approval was not necessary.

### Consent for publication

Not applicable.

### Availability of data and materials

Datasets supporting the conclusions of this article are available for non-commercial use via a data use agreement (DUA) with The Maschhoff’s, LLC.

## Authors’ contributions

JED and CJG conceptualized and designed the research. CJG managed data acquisition, storage, and retrieval. CJG estimated variance components and performed association analyses. All authors interpreted results. CJG and JED wrote the initial version of the manuscript, which was edited by all authors. All authors read and approved the final manuscript.

## Acknowledgements

We acknowledge research managers of The Maschhoff’s, LLC for collection and curation of the genotype data and meta-data. We appreciate the support by The Maschhoff’s, LLC in terms of funding and generation of data for continued collaboration. The computation for this work was performed on the high performance computing infrastructure provided by Research Computing Support Services and in part by the National Science Foundation under grant number CNS-1429294 at the University of Missouri, Columbia MO.

